# Quantifying Contributions from Topological Cycles in the Brain Network towards Cognition

**DOI:** 10.1101/2024.06.03.597217

**Authors:** Sumita Garai, Sandra Vo, Lucy Blank, Frederick Xu, Jiong Chen, Duy Duong-Tran, Yize Zhao, Li Shen

## Abstract

This study proposes a novel metric called Homological Vertex Importance Profile (H-VIP), utilizing topological data analysis tool persistent homology, to analyze human brain structural and functional connectomes. Persistent homology is a useful tool for identifying topological features such as cycles and cavities within a network. The salience of persistent homology lies in the fact that it offers a global view of the network as a whole. However, it falls short in precisely determining the relative relevance of the vertices of the network that contribute to these topological features. Our aim is to quantify the contribution of each individual vertex in the formation of homological cycles and provide insight into local connectivity. Our proposed H-VIP metric captures, quantifies, and compresses connectivity information from vertices even at multiple degrees of separation and projects back onto each vertex. Using this metric, we analyze two independent datasets: structural connectomes from the Human Connectome Project and functional connectomes from the Alzheimer’s Disease Neuroimaging Initiative. Our findings indicate a positive correlation between various cognitive measures and H-VIP, in both anatomical and functional brain networks. Our study also demonstrates that the connectivity in the frontal lobe has a higher correlation with cognitive performance compared to the whole brain network. Furthermore, the H-VIP provides us with a metric to easily locate, quantify, and visualize potentially impaired connectivity for each subject and may have applications in the context of personalized medicine for neurological diseases and disorders.

## 1 Introduction

Understanding the precise mechanisms underpinning cognitive functions of the brain has remained a long time goal of neuroscience. By addressing the question of how cognitive activities are affected or controlled by neural circuits in the brain, one can help with treatment and prevention plans for various diseases and disorders inducing cognitive malfunction such as Alzheimer’s Disease [1], Fronto-temporal Dementia [2], Parkinson’s Disease [3], Traumatic Brain Injury [4], Attention Deficit Disorder [5, 6], and other forms of cognitive impairments. Having a robust understanding of cognitive processes can also help healthy adolescents and adults with memory and learning, productivity optimization, and smooth execution of their day-to-day cognitive tasks. Modern tools such as functional neuroimaging, tractography, electrophysiology, cognitive genomics have been found to be useful in uncovering the functions and dynamics of various brain regions [7]. Network Neuroscience [8–10] is a subfield of neuroscience that uses various imaging modalities to construct graphical representations of relationship/connectivity between brain regions and analyze brain structures and associated functions based on these connectivity patterns. The human cortex is parcellated into disjoint brain regions of interest (ROIs) forming the notion of nodes or vertices. The number of ROIs can vary widely, ranging from a few dozen to several hundred. Functional and structural connectomes are constructed by quantifying the connectivity between these different ROIs.

Typically, diffusion-MRI (dMRI) is used to obtain the structural connectomes. Diffusion-MRI uses in-vivo diffusion of water molecules that are excited with the imposition of a strong magnetic field to trace the white-matter fiber tracks between grey matter regions in the brain. From these fiber tracks, structural connectivity edges connecting each pair of ROIs are often defined by measures such as length of fibers, number of fibers, fiber anisotropy, fiber density, to name a few.

Functional connectomes refer to the brain’s functional connectivity patterns, which is usually represented by the correlations of activities between different brain regions, often derived from functional magnetic resonance imaging (fMRI). The majority of functional connectome studies utilize either resting state functional magnetic resonance imaging (rs-fMRI), acquired by scanning the subject at rest, or task fMRI, which involves scanning the brain while the subject performs specific cognitive, sensory or motor tasks. Functional connectomes provide valuable insight into the dynamic and integrated nature of brain activity. Connectomics analysis can facilitate a deeper understanding of brain architecture and functionality, pinpoint specific differences between healthy brain versus various neurological and psychiatric disorders, which can further help with diagnosis, understanding disease mechanisms, and developing potential treatments [9, 11–13].

Even though connectomics analysis is one of the very few methods available for *in vivo* analysis of human brain in a non-invasive manner, it has several limitations. Some of these limitations include data quality and variability, limited spatial and temporal resolution, lack of consensus in parcellation schemes and thresholding parameters and interpretation of results [14–18]. In the past decade, topological data analysis tools such as Persistent Homology have proven to be a helpful solution for conducting threshold-free network analysis, especially in the context of brain connectomics [19–24]. Persistent Homology offers insights into the network connectivity through understanding of connected components, cycles and cavities (zero-dimensional, one-dimensional and two-dimensional persistent homology, respectively).

A fundamental problem arising in the practical applications of topological data analysis is its interpretability: given a topological feature, how can we understand its significance in terms of the underlying data? For example, a study [25] exploring connection between synaptic connectivity network and brain function showed that in response to stimuli, correlated activity binds synaptically connected neurons into functional cliques and cavities with increasing complexity, explaining the observed abundance of cliques of neurons. Another study which uses homological scaffolds reveals that significant changes occur in the brain’s network topology post-psilocybin infusion compared to a placebo [23].

To explore the connection between persistent cycles and cognition, in a previous study [26], we defined two network-level summary statistics: **average persistence** and **persistence entropy**. We found these measures to have remarkable predictive power for cognitive performance. Using only two features (average persistence and persistence entropy) for each subject, we noted a substantial correlation between the predicted cognitive performance and the actual cognitive score.

While summary statistics are useful for understanding the overall trends, they do not provide insights into mesoscopic connectivity patterns or the *individual contributions* of each brain region to various cognitive tasks.

In this study, we address the aforementioned question by quantifying local contribution from each region. With this goal in mind, we developed a new metric which can help with refined understanding of network topology, not only globally but also locally, by assigning a score to each region of interest. We call this metric Homological Vertex Importance Profile (H-VIP) of the subject, which gives a vector of length equal to the size of the network, with each component representing the significance score of each ROI the network. We note that traditional network measures (cf. Brain Connectivity Toolbox, [10]), such as betweenness centrality, participation coefficient, and nodal strength, offer tools for assessing the role of each vertex in facilitating network connectivity. However, it is not easy to estimate how these measures affect the *global* topology of the network, particularly regarding topological cycles. The H-VIP metric proposed here is an attempt to provide a top-down or global-to-local measure for understanding topological cycles in a network. In essence, this metric captures and compresses connectivity information from nodes which are at *multiple degrees of separation* and and projects back onto local nodes.

We apply this metric to two different data-sets: (1) whole brain structural network and the frontal sub-network for subjects (young healthy adults) from the Human Connectome Project and (2) functional network extracted using fMRI from the Alzheimer’s Disease Neuroimaging Initiative Database. We use our propsed H-VIP metric to predict the cognitive scores of the participants by applying a linear regression model. We compute the correlation between the predicted cognitive scores and the actual cognitive scores and find that there is a statistically significant correlation between the predicted scores and the true scores.

Our findings adds to the evidence in the support of the hypothesis that there is indeed an association between topological features of the brain network and cognitive measures such as fluid intelligence. In our analysis, we also find that H-VIP metric has prediction power for the cognitive decline in Alzheimer’s disease. The Vertex Importance Profile captures individual differences in the brain network, as well as assigns significance scores (or ranks) to each ROI’s contribution to each subjects overall connectivity profile. Understanding the role of each brain region and, more importantly, being able to *quantify* their role with respect to a given neuropathology would greatly help us understand the probable mechanism of the disease/disorder and design an individualized medical intervention targeted to fit each person’s profile.

## 2 Methods and Materials

### 2.1 Data and Preprocessing

Data were downloaded and analyzed under approval of the University of Pennsylvania Institutional Review Board.

#### 2.1.1 Human Connectome Project Cohort

The Human Connectome Project for Young Adults (HCP-YA) is an initiative to provide a detailed and comprehensive understanding of the organization of the brain’s network during early adulthood, a critical period of brain maturation and development where many patterns and changes in brain connectivity could occur. The Human Connectome Project has enabled researchers to investigate how brain connectivity is related to cognitive abilities, decision making, emotional processing, and other aspects of human behavior in young adults by detailed comparisons between brain circuits, behaviors, and genetics at the individual subject level [27]. For more information, please refer to www.humanconnectome.org.

In this study, we accessed the prepossessed imaging data through the ConnectomeDB, and the structural connectomes are extracted using the pipeline described in [28]. The Desikan-Killiany (DK) atlas [29] is used to define the ROIs corresponding to the vertices in the structural connectome with 68 cortical surface regions (34 vertices in each hemisphere) and 19 subcortical regions. For each pair of ROIs, the streamlines connecting them are extracted using the following procedure:

- Each gray matter ROI is dilated to include a small portion of white matter regions.
- Streamlines connecting multiple ROIs are cut into pieces in an attempt to extract the correct and complete pathway.
- Outlier streamlines are removed.

In total, 1,065 brain structural connectomes are extracted from the latest release of the HCP-YA dataset. From this dataset we further separate out 399 subjects which are neither twins nor siblings using the demographic data on parental identifiers. In this paper, we performed our analysis on the aforementioned subset of subjects. Please refer to Table 1 for a quick overview of the demographics of the participants and the cognitive scores relevant to our study. This study was approved by the institutional review boards of all participating institutions, and written informed consent was obtained from all participants or authorized representatives.

**Table 1:** Subject demographics and Cognitive Measures of the 399 subjects (214 male/185 female) from the HCP-YA cohort.

| Attributes | Range | Mean ± Stand. dev. |
| --- | --- | --- |
| Age (in years) | 22 – 36 | 28.51 ± 3.69 |
| Penn Matrix Reasoning Test (PMAT24) | 06.00 – 24.00 | 17.24 ± 4.69 |
| Oral Comprehension | 86.20 – 150.70 | 116.77 ± 10.30 |
| Oral Comprehension, age-adjusted | 65.02 – 138.08 | 106.87 ± 14.46 |
| Picture Vocabulary Test | 92.84 – 153.08 | 116.85 ± 9.19 |
| Picture Vocabulary Test, age-adjusted | 69.45 – 153.08 | 109.38 ± 14.56 |
| Computerized Penn Word Memory Test | 26.00 – 40.00 | 35.88 ± 2.79 |
| Pattern Composite Processing Speed | 60.09 – 154.69 | 115.06 ± 15.96 |

#### 2.1.2 Alzheimer’s Disease Neuroimaging Initiative Cohort

Data used in the preparation of this article were obtained from the Alzheimer’s Disease Neuroimaging Initiative (ADNI) database (adni.loni.usc.edu) [30, 31]. The ADNI was launched in 2003 as a public-private partnership led by principal investigator Michael W. Weiner, MD. The primary goal of ADNI has been to test whether serial MRI, PET, other biological markers, and clinical and neuropsychological assessment can be combined to measure the progression of early cognitive impairment leading to dementia. All participants provided written informed consent and study protocols were approved by each participating site’s Institutional Review Board (IRB). Up-to-date information about the ADNI is available at www.adni-info.org.

From the latest ADNI-3 data release, we studied functional MRI scans of 406 subjects. These images were processed in-house, using the Python package Connectome Mapper [32]. This cohort consists of three different subcategories of subjects: cognitive normal (CN), mild cognitive impairment (MCI), and dementia (AD). Lausanne [33] atlas with 99 regions of interests (ROIs) was used as the parcellation scheme. Table 2 provides an overview of the characteristics of the participants such as age, gender, education with breakdown based on disease-pathology, as well as the cognitive markers investigated in this study.

**Table 2:**
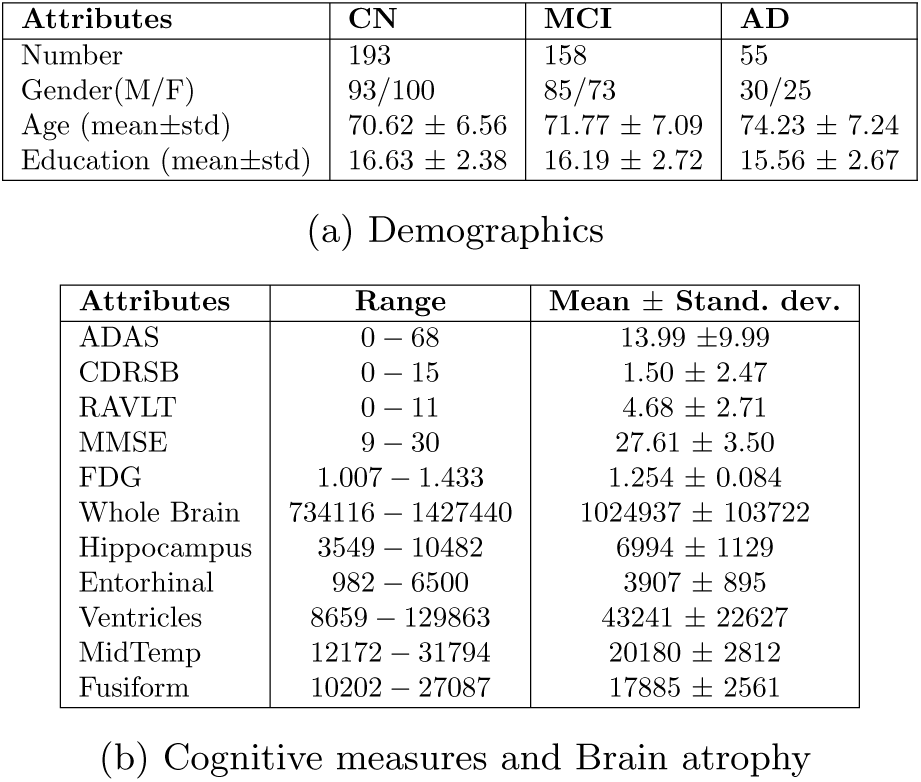
Subject demographics for ADNI cohort.

### 2.2 Methods

#### 2.2.1 Persistence Barcode

A brain network can be represented as a weighted graph *G* = (*V, w*) comprising of a set of vertices (or nodes) *V* and edge weights *w* = (*w_ij_*) given by real numbers. The vertices *V* correspond to brain ROIs and weights *w* = (*w_ij_*) represents strength of the connections. The number of vertices of *G* is given by *|V |* = *n*. Note that a complete graph on *n* vertices has *n*(*n −* 1)*/*2 maximum possible number of edges.

A binary graph *G_ϵ_* = (*V, w_ϵ_*) of *G* is defined as a graph consisting of the node set *V* and binary edge weight *w_ϵ,ij_* = 1 if *w_ij_ > ϵ* or 0 otherwise. The binary graph network *G_ϵ_* is the 1-skeleton of the simplicial complex which consists solely of these vertices and edges. We compute the homological groups, informally referred to as holes, of the simplicial complex associated to *G_ϵ_*. In such 1-skeleton, 0-dimensional (0D) homological groups represent *connected components* and 1-dimensional (1D) homological groups represent *cycles*. The rank of the zero dimensional and the one dimensional homological group, which also represents the number of connected components and cycles, are referred to as the 0*^th^* Betti number *β*_0_(*G_ϵ_*) and the 1*^st^* Betti number *β*_1_(*G_ϵ_*), respectively. A graph filtration of *G* is defined as a collection of nested binarized networks:

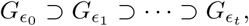

where *ϵ*_0_ *< ϵ*_1_ *< · · · < ϵ_t_* are filtration values, often lies in the interval [0, 1] for weight-normalized graph *G*. With increasing filtration value *ϵ*, lower connectivity edges are removed resulting in a sparser network and a change in the 0*^th^* and 1*^st^* homology groups and the corresponding Betti numbers.

Persistent homology keeps track of appearance (birth) and disappearance (death) of connected components and cycles over filtration values *ϵ*, by keeping track of the aforementioned Betti numbers. A set of intervals 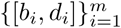is associated to keep track of birth and death of a *persistence*. This set of intervals is referred to as Persistence Barcode [20]. Lifetime, *l_i_* = (*d_i_ − b_i_*) of a persistence signifies that at the filtration value *b_i_*, a connected component or a cycle appeared; and at filtration threshold *d_i_*, a component or cycle disappeared or died. Long persistence is believed to signify topologically relevant features whereas short persistence indicate noise [34]. A detailed discussion on the mathematical foundation of homology and co-homology [35] and several relevant studies on applications of persistence barcodes can be be found in the literature [19, 34, 36–38].

#### 2.2.2 Homological Vertex Importance Profile

In a previous study [26], we defined two whole-brain topological measures: **average persistence** and **persistence entropy**. By utilizing these two features (average persistence and persistence entropy) for each subject, we observed a significant correlation between the predicted cognitive performance and the actual cognitive score. Although these global network measures are valuable, they do not offer insights into the specific local connectivity patterns or the unique contributions of individual brain regions to various cognitive tasks. To complement our previous work, here we present a novel metric to quantify the importance of each node of the network in forming cycles.

To define this metric, firstly we need to find representative cycles that captures the structure of each cycle in terms of vertices, and we use co-homology for identifying this representation. Mathematically, co-homology is the *dual* of homology. Since computing homology groups is a difficult task, it is often computationally convenient to work with co-homology and co-cycles instead, and in this article, we will refer to them interchangeably. Another challenge is that representative co-cycles are not unique. For a given co-cycle class, we have several representative co-cycles. Each representative co-cycle of a given equivalence class (also known as Chern class, which are unique up to contractible sub-spaces) generate the same cohomology class. We overcome this challenge by meticulously keeping track of all representatives, as there are only finitely many of them.

In this study, we use the Python package Ripser [39] to compute the representative co-cycles. Once we compute the 1D cycles and the corresponding representative cocycles for the brain network or sub-network, we can use these representatives to define a metric: **Vertex Importance (VI)** for any given vertex by quantifying its contribution to all the 1D-persistence (or cycles) occurring in this network. The **Homological Vertex Importance Profile** or **H-VIP** of a network is a vector consisting of relative importance of each vertex of the network in forming 1D co-cycles. Formally,

**Definition 1.** *Let G be a graph network with n vertices and* 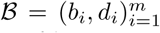 *be its associated 1D persistence barcode. If* 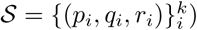 *is the set of k representatives corresponding to a particular co-cycle 𝒞, then the Vertex Importance of the vertex v will be the average number of times it appears in all the representatives for 𝒞, i.e*.

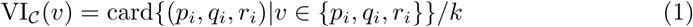

Note that the metric vertex importance VI*_𝒞_*(*v*) depends on the chosen cycle *C* and the vertex *v*.

**Definition 2.** *The Homological Vertex Importance Profile of the network G is given by the tuple:*

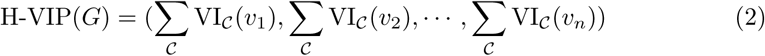

*where indices in the sum runs over all the co-cycles 𝒞 of the network G*.

This profile gives an estimate for the significance of the contribution (in forming cycles) of a particular vertex. Note that this is a vector of dimension *n*, the number of vertices in the graph.

Please refer to the schematic diagram in Figure 2 for a visual intuitive comparison of the H-VIP and the graph-theoretic local measures: Degree Centrality (DC) and Betweenness Centrality (BC) [10]. This diagram illustrates a graph where the vertex colored red *v*_4_ demonstrates the highest vertex importance, indicating its involvement in the greatest number of homological cycles. Meanwhile, the vertex *v*_6_ (colored green) is positioned in-between maximum number of shortest paths, enjoying maximum betweenness centrality. Additionally, the vertex in blue *v*_7_ showcases the highest degree of centrality or nodal strength. We note that DC captures connectivity information at only one degree of separation. While BC does take into account of all the shortest paths that the given vertex lies on, it does not account for the closed loops or cycles. Significance of a cycle lies in the fact that all the vertices forming a cycle communicate with each other, *albeit* indirectly.

**Fig. 1:**
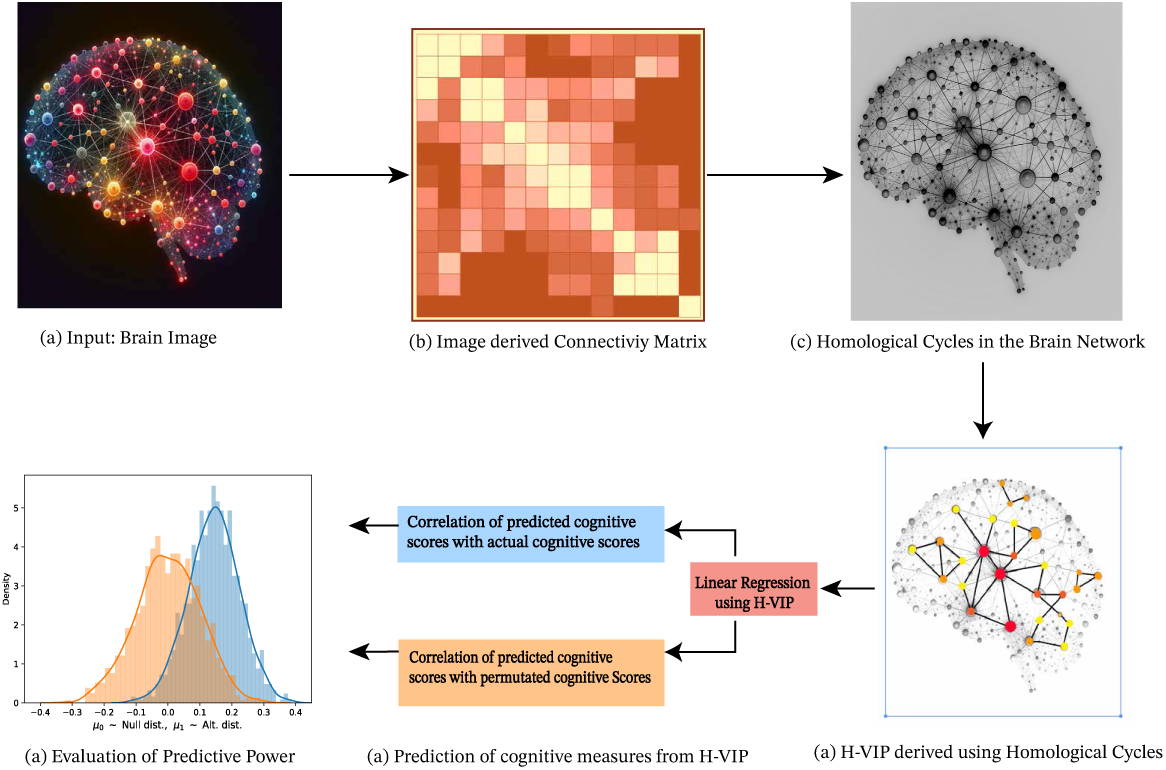
Workflow: Predicting of Cognitive score using Vertex Importance Profile

**Fig. 2:**
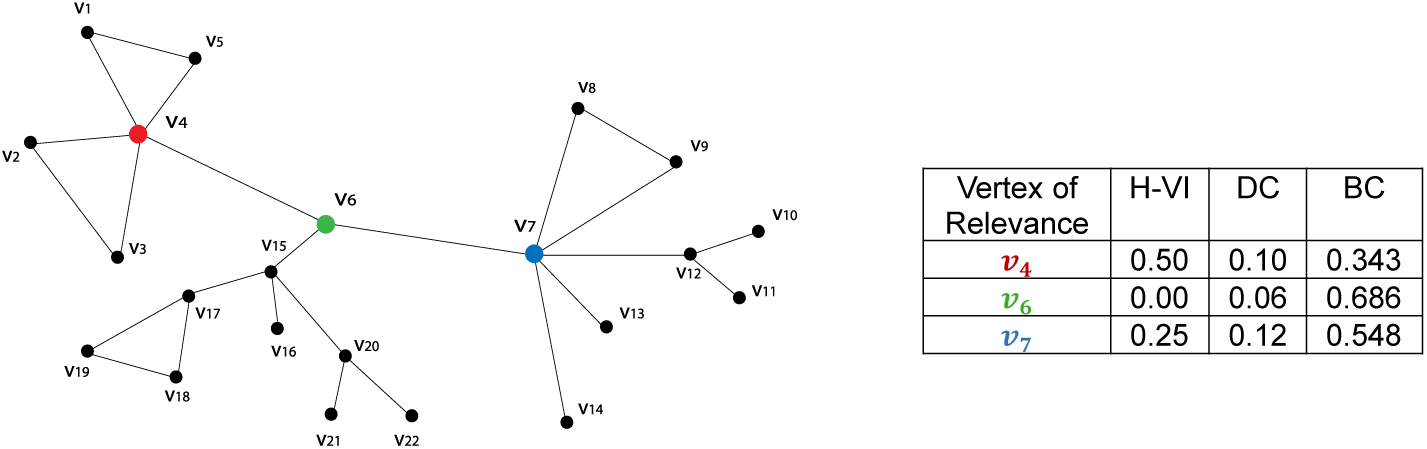
Vertex *v*_4_ (highlighted in red) has highest H-VI as it is contributing to maximum number of cycles. Vertex *v*_6_ (green) has highest Betweenness Centrality and breaking it would disconnect most of the shortest paths. Vertex *v*_7_ (blue) has highest degree centrality or nodal strength.

We apply this new metric to extract the homological vertex importance profile vector for the whole brain network as well as the frontal sub-network for each participant in our study. Next we use these vectors for predicting the cognitive scores of the participants, by applying a linear regression model.

#### 2.2.3 Prediction using H-VIP

First, we split our subjects into two categories: training and testing, in the ratio 80% and 20% respectively, chosen at random. For each cognitive measure, we use our feature vector H-VIP to train our linear regression model with the training subjects (80%). The we use the rest of the 20% of subjects (test subjects) to predict the cognitive score and calculate the Pearson correlation between the predicted value of the score and the actual score. The performance is reported on the testing set only, so there is no overfitting bias. We repeat this process of splitting data into training and testing subjects 1000 times, each time with a different randomization seed. Each iteration gives us a correlation value, which results in a normal distribution with mean *µ*_1_, shown in color blue, see Figure 3 and Fig 6. Next, we permute/shuffle (using a randomization seed) the cognitive scores of the 20% test subjects and repeat the above process i.e. we calculate the Pearson correlation between the randomly assigned score and the predicted score, which again gives us another normal distribution, with mean *µ*_0_, shown in color orange in Figure 3 and Fig 6. Under the null hypothesis we expect no statistically significant difference between these two distributions. In the next section, we summarize our findings.

**Fig. 3:**
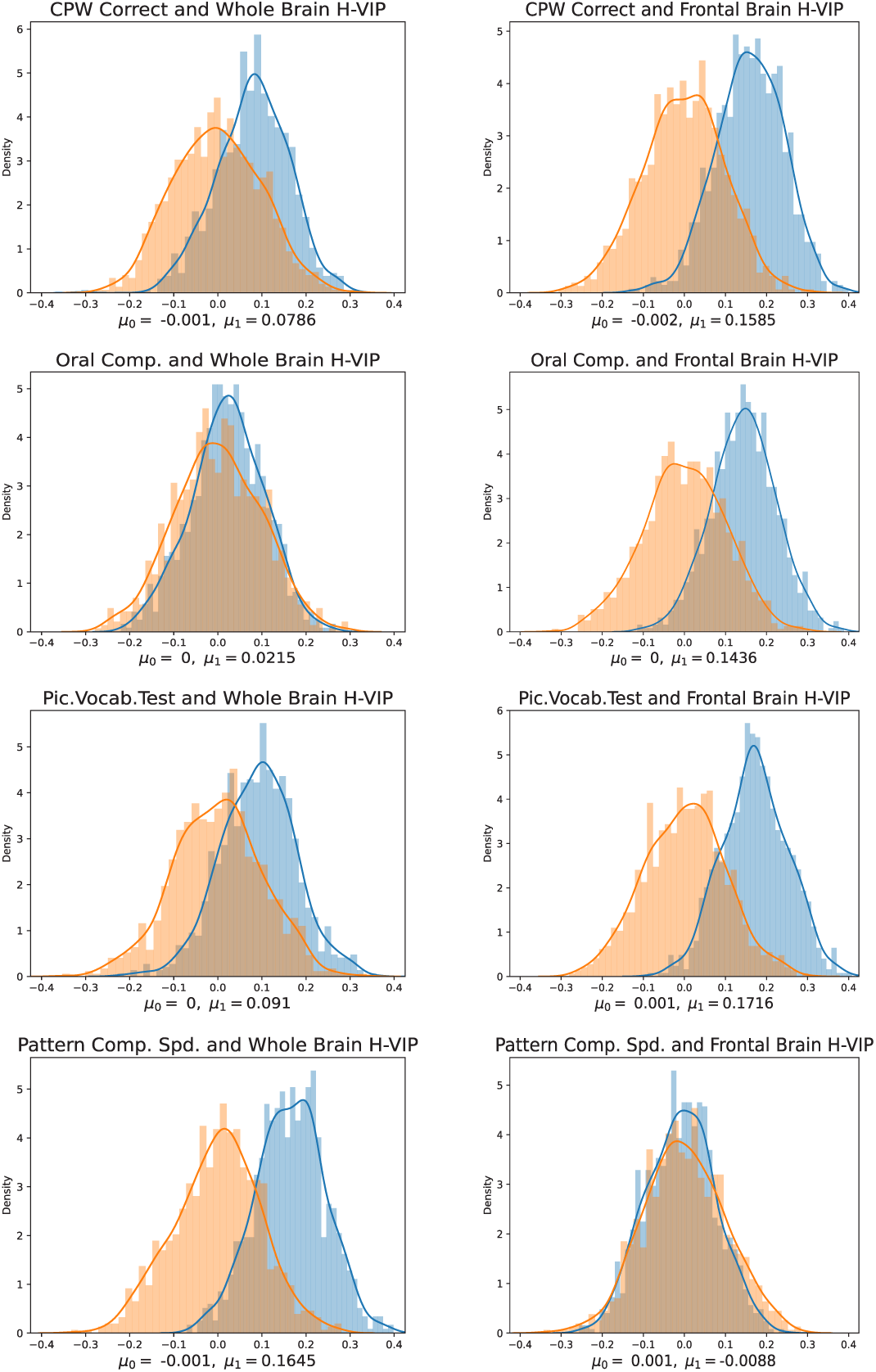
Correlation between cognitive scores predicted by H-VIP and the true cognitive scores (show in blue), plotted against correlation between scores predicted by H-VIP and randomly assigned scores (shown in orange).

## 3 Results

### 3.1 Study of cognitive scores from the HCP-YA subjects

For the HCP dataset, we performed regression using the following set of features:

- Homological Vertex importance profile for the Whole Brain
- Homological Vertex importance profile for the Frontal Brain sub-network.

We observe in Figure 3 that correlation between the cognitive score predicted by the H-VIP and the shuffled (and randomly assigned) cognitive scores forms a normal distribution with mean approximately zero, showing evidence that randomly assigned scores are not correlated with the predicted scores. However, we observe *a right shift* or positive correlation between the predicted scores and the *actual* cognitive scores. We also compare network measures DC and BC to our proposed measure H-VIP to demonstrate its potential, see Table 3, where statistically significant differences are highlighted in bold.

**Table 3:**
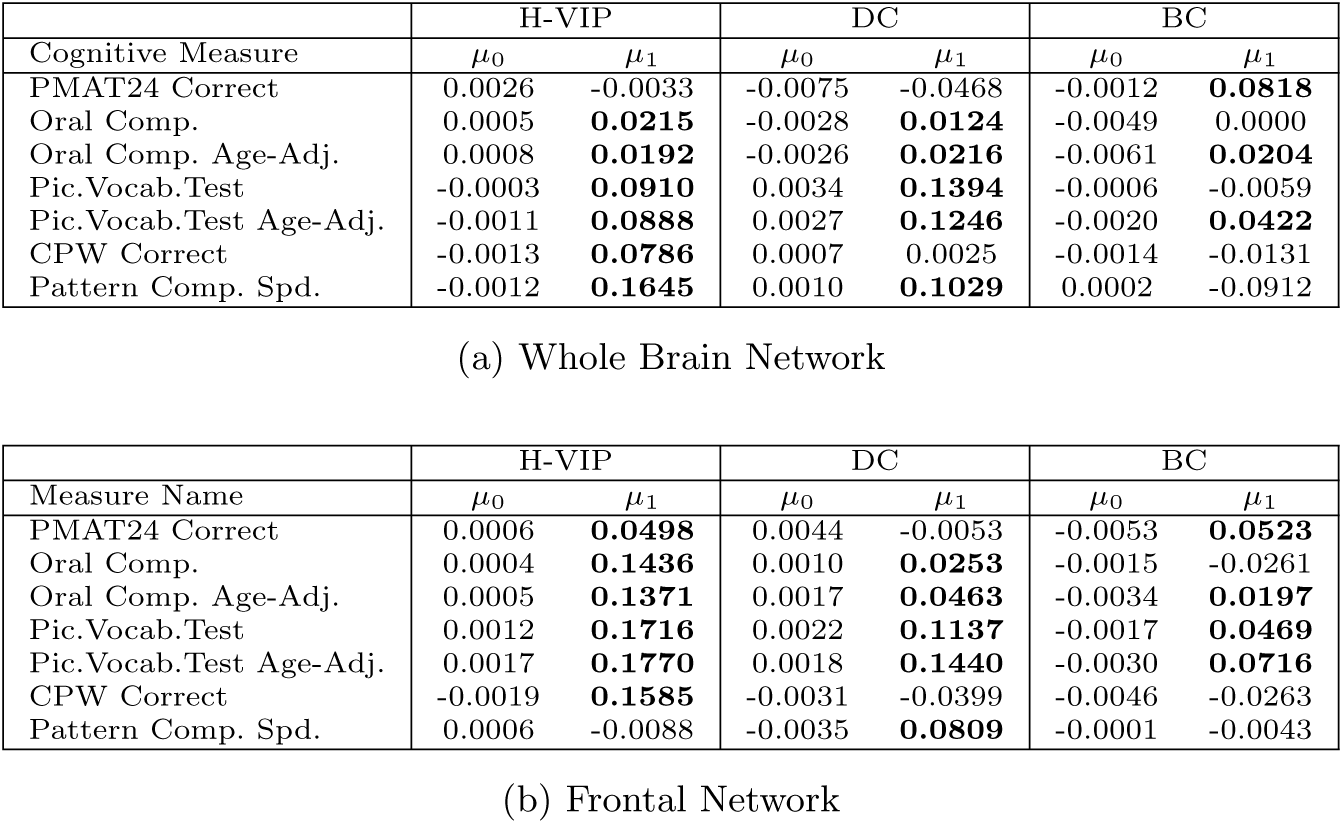
Correlation of network measures with Cognitive Measures.

Interestingly, we observe that H-VIP and DC of the frontal lobe sub-network comprising of 20 regions of interest (ROIs) has a significantly higher correlation with cognitive performance compared to the whole brain network. This provides evidence that there is indeed a distinctive topological structure underlying the frontal brain network correlated to cognitive performance, and corroborates with the existing literature [40–42]. We note that H-VIP is a better predictor of cognitive performance for most of the cognitive measures considered in this study.

The Figure 4 shows the average (taking into account all subjects) H-VIP for the subjects from HCP-YA under this study. This average H-VIP conveys a fairly simple yet reasonably accurate representation of the degree of involvement of each node in the frontal area of a healthy adult brain of population of age between 22-35. In Figure 5, we see the individualized H-VIP, three subjects ranked minimum, median (50 percentile) and maximum on their age-adjusted picture vocabulary scores, and we do indeed see striking differences in the H-VIP for the minimum and maximum score. For a detailed discussion and possible explanation, please refer to Sec. 4.

**Fig. 4:**
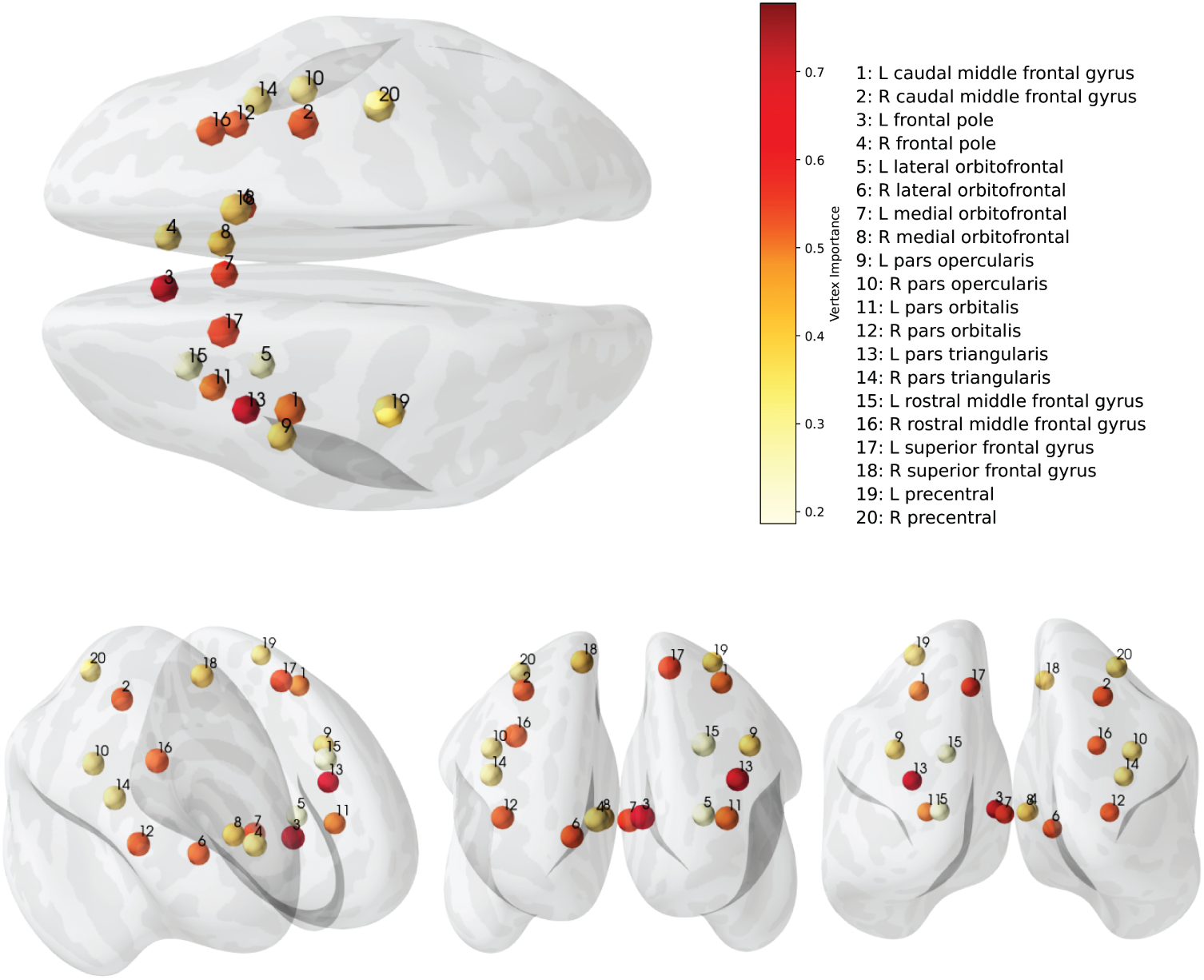
Average Homological Vertex Importance Profile for HCP-YA cohort in this study. Top: Dorsal View. Bottom: Frontal, Rostral and Caudal view from left to right assessment tests and physiological biomarkers such as beta-amyloid, tau, phospho-tau and atrophy in various brain regions, including the whole brain.

**Fig. 5:**
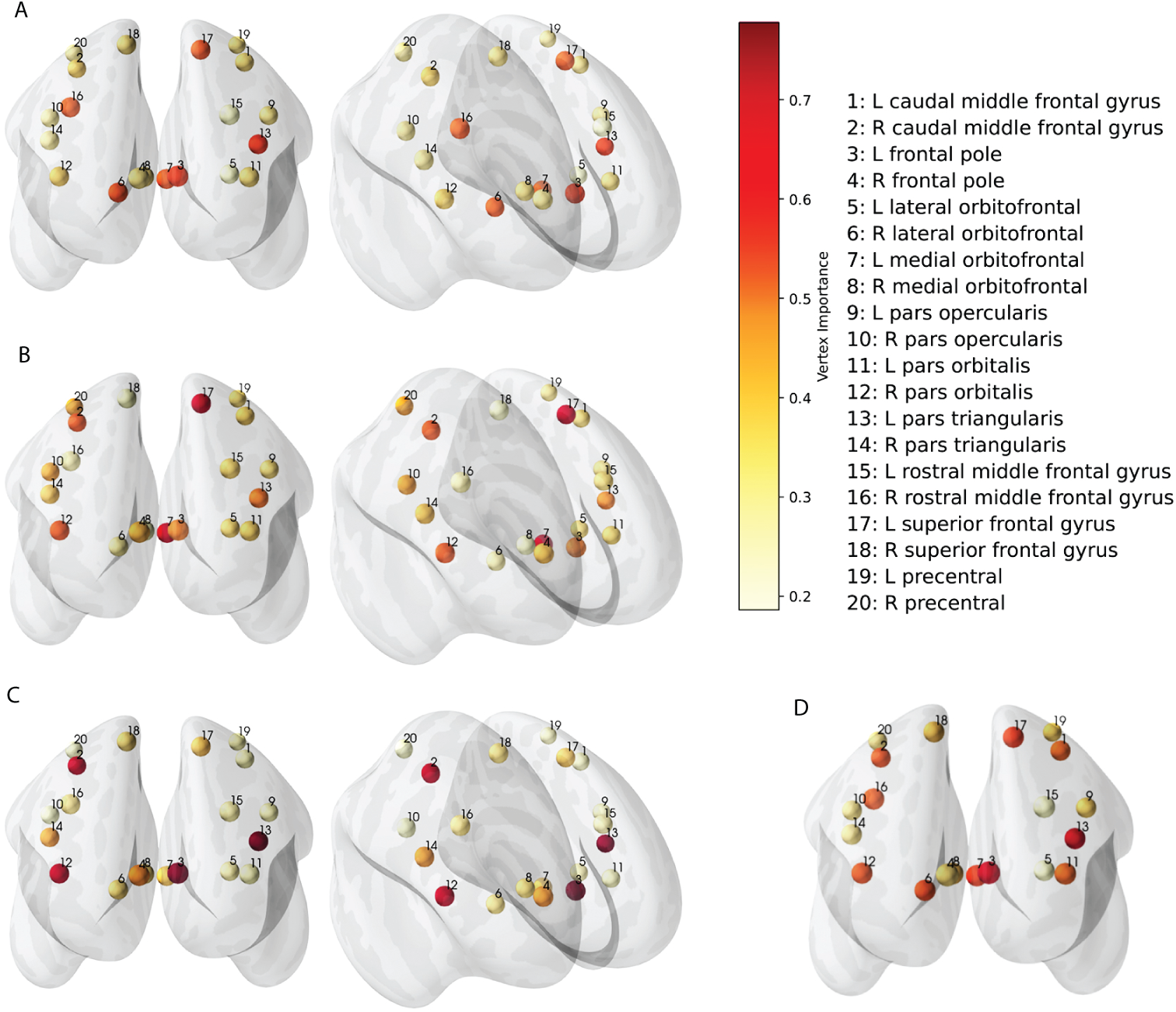
Homological Vertex Importance Profiles for subject with minimum (Fig. A), median (Fig. B) and maximum (Fig. C) Picture Vocabulary Cognitive Scores. Fig. D reproduces the average H-VIP of the study cohort for comparison.

There are several immediate applications of this profile. It can be used as standard for comparison with respect to age related brain connectivity changes, or assess cognitive neuro-diversity or neurological disorder. Also, it can further aid in discovery of distinctive brain circuitry that contribute to different types of cognitive tasks.

#### 3.1.1 Study of cognitive assessment, biomarkers and brain atrophy in Alzheimer’s disease using ADNI-3

Our next study is on a cohort of 406 subjects, ranging from healthy subjects to various levels of cognitive impairments including Alzheimer’s disease. Alzheimer’s and other neurodegenerative diseases are hypothesized to be structural disruptions that lead to functional impairments [43, 44]. We used the functional connectomes from ADNI-3 and extract the aforementioned metric H-VIP for each subject. Similar to the first dataset, we use a linear regression model to predict a set of outcomes used to characterize state or progression of Alzheimer’s diseases. These outcomes include a set of cognitive

We observe that wholebrain H-VIP has prediction power for the scores obtained from Alzheimer’s Disease Assessment Scale (ADAS) and Mini-Mental State Examination (MMSE), see Figure 6 and Table 4. The highest predictive power is for atrophy in the hippocampus and medial temporal lobe, where we observe the maximum right shift from the null distribution. The mid temporal region is crucial for episodic and spatial memory with distinct functions such as encoding, consolidation and retrieval of memory [45].

**Fig. 6:**
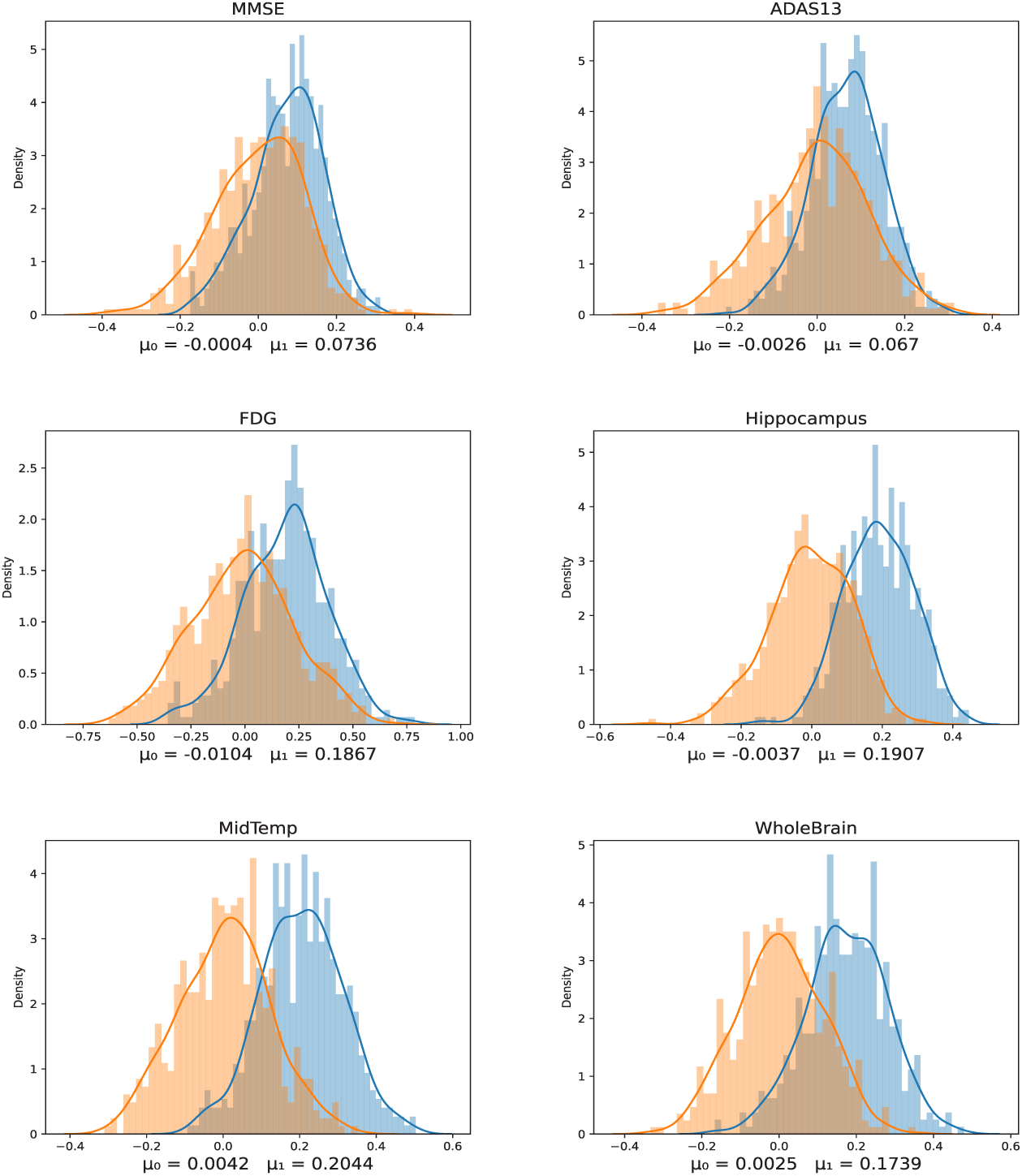
Correlation between predicted and true values of cognitive scores, biomarkers, atrophy (shown in blue), plotted against the null distribution of randomly permuted scores (shown in orange).

**Table 4:** Correlation between predicted measure using H-VIP vs actual score for the ADNI cohort.

| Measure | $\mu_0$ | $\mu_1$ |
| --- | --- | --- |
| ADAS13 | 0.0067 | **0.0670** |
| CDRSB | 0.0041 | 0.0089 |
| RAVLT | 0.0020 | 0.0504 |
| MMSE | 0.0039 | **0.0736** |
| FDG | 0.0010 | **0.1867** |
| WholeBrain | 0.0011 | **0.1739** |
| Hippocampus | 0.0074 | **0.1907** |
| Entorhinal | 0.0035 | **0.1059** |
| Ventricles | 0.0002 | 0.0152 |
| MidTemp | -0.0023 | **0.2044** |
| Fusiform | -0.0050 | **0.0858** |

For illustrations, we also plot the frontal subnetwork H-VIP of the subject with the lowest MMSE score (subject A) vs subject with highest MMSE score (subject B) from our study cohort in Figure 7. Note that subject B is cognitively normal whereas subject A has Alzheimer’s disease. Here, the H-VIP allows us to zoom into the frontal sub-network of the subjects. In Fig 7, we see that the subject with lowest MMSE score in the cohort, has relatively diminished vertex importance for most regions, *except* for the remarkably high contribution from right superior frontal region, labelled as region 9 in Figure 7A.

**Fig. 7:**
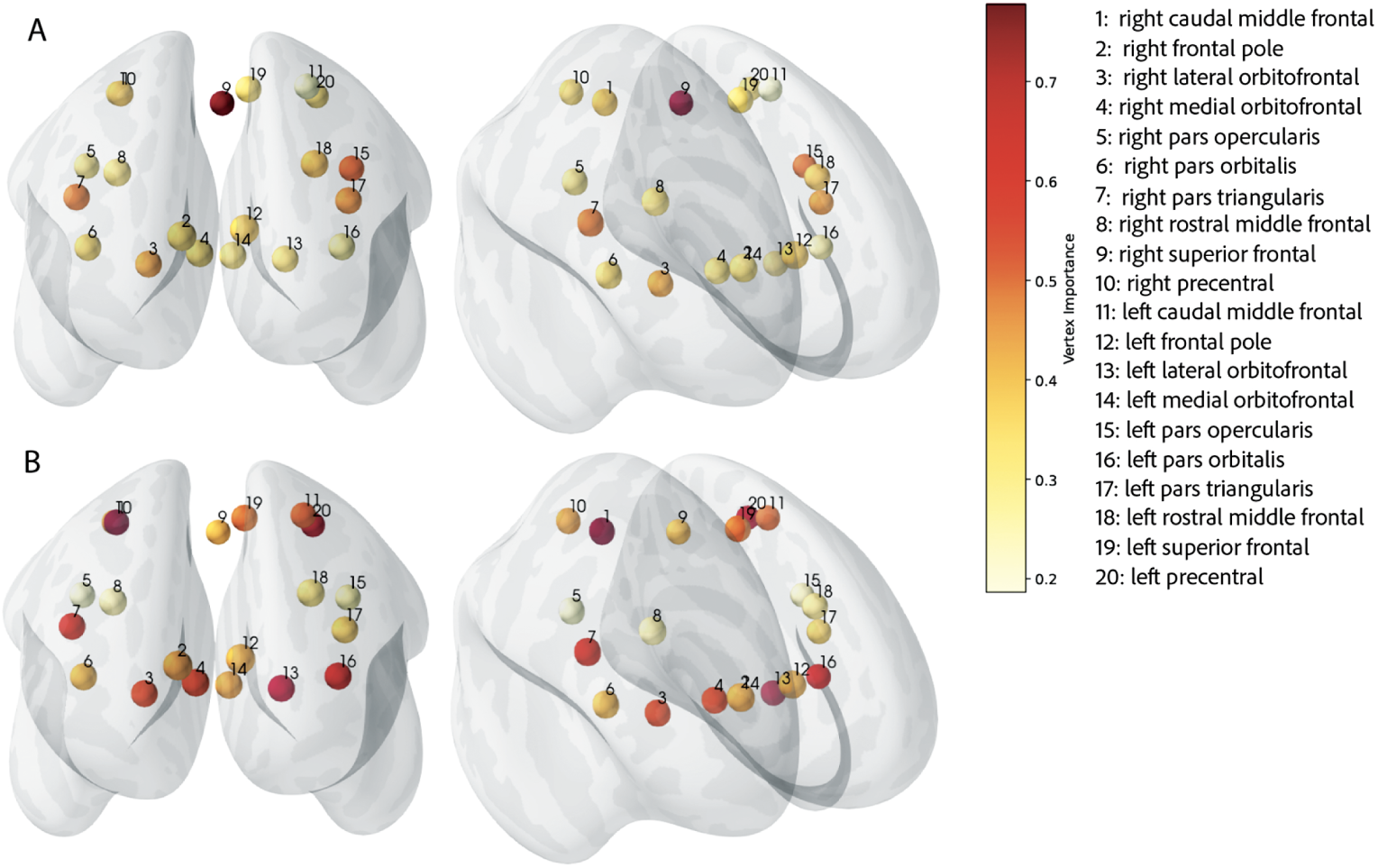
Homological Vertex importance profile of the subject with lowest MMSE score (Fig. A) and highest MMSE score (Fig. B) in the ADNI study cohort

## 4 Discussion

The goal of this study is two fold: to understand and quantify the role (or importance) of each brain region for cognitive tasks in healthy adults and to pinpoint individual anatomical differences in the manifestation of the disease pathology, such as in Alzheimer’s disease.

Anatomical brain networks are, in some sense, the true skeleton of brain architecture. Connectivity measurement tools have long been used to investigate network efficiency and clustering as means to quantify integration and segregation properties in brain networks at multiple scales [10, 46, 47]. Since efficient connectivity of a network ensures smooth transmission of information, it has been hypothesised that it is also related with fluid, crystallized, general intelligence [47–50]. To understand underlying mechanism behind behind information processing and cognitive performance, we apply our metric of vertex importance to quantify role of various regions of interest. In Fig 4 we note that, *on an average*, the regions in the left frontal pole, left superior frontal gyrus, left pars triangularis and left medial orbitofrontal cortex have higher vertex importance. The frontal pole and superior frontal gyrus are two important brain regions located in the frontal lobe of the cerebral cortex. These regions play critical roles in various aspects of cognition, which involve higher-order thinking, decision-making, and goal-directed behavior [51–54]. The medial orbitofrontal cortex (located in the frontal lobes) region has been associated with various cognitive and emotional processes, including decision-making, social behavior, emotion regulation, and reward processing [55]. It plays a key role in integrating sensory information, emotional responses, and cognitive functions to guide behavior and decision-making. The pars triangularis is one of the three subdivisions of the broader region known as the Broca’s area, which is traditionally associated with language production and comprehension [56], but also implicated in working memory and cognitive control such as regulating attention, inhibiting irrelevant information, and switching between tasks [5, 57].

Our study provides evidence that H-VIP can prove to be a valuable tool in understanding the connectivity profile of a network, as it assigns a score to each vertex (node) in the network, reflecting its significance or contribution to network dynamics. When comparing brain networks of two subjects, more often than not, we observe different H-VIPs, which effectively capture their individual differences. These differences may arise due to variations in brain structure, function, or connectivity patterns. Individual differences can be attributed to factors such as genetics, age, cognitive abilities, or the presence of neurological conditions. The significance of observing different H-VIPs between subjects lies in the potential to uncover unique patterns associated with specific cognitive functions, behaviors, or disease states. By linking these individual differences to specific network features, researchers can gain insights into how brain networks are organized and how they contribute to inter-individual variations in behavior and cognition.

Furthermore, H-VIP is relevant in the context of personalized medicine and neuroscience. They can help identify brain regions that might be more susceptible to dysfunction or alterations in neurological disorders. This knowledge can be leveraged to develop personalized treatment strategies tailored to an individual’s specific neurobiological characteristics. Overall, the H-VIP has potential to be a powerful tool in network neuroscience, shedding light on individual differences in brain connectivity and offering refined understanding of nuances in the progression of various neurological and psychiatric conditions.

There are certain limitations inherent to any study of this kind. The estimation of whole-brain structural connectivity is notably influenced by data processing parameters, such as the number of streamlines used, different parcellation schemes, diverse data processing pipelines, and varying resolutions. With due caution, we are still able to extrapolate behavioral metrics from the anatomical connectivity networks, keeping in mind our proposed topological characteristics are sensitive to the aforementioned parameters. The functional connectivity network is slightly more complex to interpret, because it is built from co-activation of regions measured from blood-oxygen-level-dependent signals, and have no *direct* underlying anatomical framework. Despite this constraint, functional networks can effectively monitor brain activity areas and distinguish areas specific to certain categories of tasks. A relevant future application of H-VIP will be to analyze various task-based (functional) networks, and isolating regions of interests activated during specific tasks.

## 5 Conclusions

In this work, we have introduced a new metric: Homological Vertex Importance Profile, which provides a mathematically rigorous formulation of the abstract yet intuitive notion of degree of participation from individual vertices in forming homological cycles in a graph network. This formulation of H-VIP provides a global-to-local projection of homological relevance of the vertices and can be thought of as an encapsulation of quasi-local connectivity profile of the network.

We have applied this metric to analyze two data-sets: structural connectomes from the Human Connectome Project for Young Adults (HCP-YA) and functional connectomes from ADNI-3. In both cases we have used H-VIP to predict cognitive performance and found that there is a positive correlation between various measures of intelligence and cycles in the anatomical and functional brain network, demonstrating its prediction power.

Our study provides evidence that higher degree of participation in forming cycles in the structural as well as functional connectivity networks in the the brain is correlated with better cognitive abilities. It also suggests that frontal lobe structural connectivity has a higher contribution towards cognitive performance relative to whole brain structural network. We conclude with the remark that H-VIP is a powerful metric that can be applied to any general network to infer interesting properties of connectivity, and certainly not limited to brain connectomes.

## Data and Code Availability

The source code and documentations used for the computation of vertex importance profile are provided in the repository https://github.com/sumitagarai/H-VIP. The authors welcome requests for additional information or assistance regarding the code and model. Both data-sets (HCP and ADNI-3) used in this study are publicly available.

## Author Contributions

S.G.: Conceptualization; Coding - Implementing the Method and application on the HCP Dataset; Writing - Original Draft, Methodology, Visualization. S.V.: Coding – Brain Visualization. L.B.: Coding - Replicating the H-VIP computation on ADNI dataset. F.X.: Coding - ADNI Data preprocessing and preparation. J.C.: Coding - ADNI Data preprocessing and preparation. D.D.-T.: Writing - Review & Editing. Y.Z.: HCP Data preprocessing and preparation, Supervision. L.S.: Conceptualization, Supervision, Methodology, Writing- Review & Editing.

## Acknowledgments

Data collection and sharing for this project was funded by the Human Connectome Project, WU-Minn Consortium (Principal Investigators: David Van Essen and Kamil

Ugurbil; 1U54MH09-1657) funded by the 16 NIH Institutes and Centers that support the NIH Blueprint for Neuroscience Research; and by the McDonnell Center for Systems Neuroscience at Washington University.

Data used in preparation of this article were obtained from the Alzheimer’s Disease Neuroimaging Initiative (ADNI) database (adni.loni.usc.edu). As such, the investigators within the ADNI contributed to the design and implementation of ADNI and/or provided data but did not participate in analysis or writing of this report. A complete listing of ADNI investigators can be found at: http://adni.loni.usc.edu/wp-content/uploads/how_to_apply/ADNI_Acknowledgement_List.pdf.

Data collection and sharing for this project was also funded by the Alzheimer’s Disease Neuroimaging Initiative (ADNI) (National Institutes of Health Grant U01 AG024904) and DOD ADNI (Department of Defense award number W81XWH-12-2-0012). ADNI is funded by the National Institute on Aging, the National Institute of Biomedical Imaging and Bioengineering, and through generous contributions from the following: AbbVie, Alzheimer’s Association; Alzheimer’s Drug Discovery Foundation; Araclon Biotech; BioClinica, Inc.; Biogen; Bristol-Myers Squibb Company; CereSpir, Inc.; Cogstate; Eisai Inc.; Elan Pharmaceuticals, Inc.; Eli Lilly and Company; EuroImmun; F. Hoffmann-La Roche Ltd and its affiliated company Genentech, Inc.; Fujirebio; GE Healthcare; IXICO Ltd.; Janssen Alzheimer Immunotherapy Research & Development, LLC.; Johnson & Johnson Pharmaceutical Research & Development LLC.; Lumosity; Lundbeck; Merck & Co., Inc.; Meso Scale Diagnostics, LLC.; NeuroRx Research; Neurotrack Technologies; Novartis Pharmaceuticals Corporation; Pfizer Inc.; Piramal Imaging; Servier; Takeda Pharmaceutical Company; and Transition Therapeutics. The Canadian Institutes of Health Research is providing funds to support ADNI clinical sites in Canada. Private sector contributions are facilitated by the Foundation for the National Institutes of Health (www.fnih.org). The grantee organization is the Northern California Institute for Research and Education, and the study is coordinated by the Alzheimer’s Therapeutic Research Institute at the University of Southern California. ADNI data are disseminated by the Laboratory for Neuro Imaging at the University of Southern California.

